# Nutrient-dependent pathology in mitochondrial hypertrophic cardiomyopathy model

**DOI:** 10.1101/2025.09.24.678132

**Authors:** Swagat Pradhan, Nahid A Khan, Tuula Manninen, Aleksandra Zhaivoron, Anu Suomalainen

## Abstract

**Objective:** Mitochondrial translation defects are a major cause of early childhood hypertrophic cardiomyopathy (CMP). While the genetic basis of these disorders is being increasingly uncovered, the downstream molecular mechanisms driving disease pathogenesis remain poorly understood. In this study, we investigated the consequences of defects in mitochondrial ribosomal large subunit protein 44 (MRPL44), associated with infantile-onset CMP in human cardiomyocytes, in nutrient environments relevant to cardiac development.

**Methods:** Induced pluripotent stem cell line with MRPL44 patient mutation and controls were differentiated to cardiomyocytes and grown in glucose or lipid-enriched medium reflecting prenatal or postnatal fuel preferences, respectively. Mitochondrial, lipid metabolic and cellular characteristics were studied by immunofluorescence and cellular transcriptome by RNA-sequencing.

**Results:** In glucose-rich medium, patient-derived cardiomyocytes exhibit increased mitochondrial DNA (mtDNA) content and elevated mitochondrial transcripts. In contrast the lipid-enriched medium triggered both mitochondrial and endoplasmic reticulum -related stress responses, disrupted lipid and cholesterol homeostasis, accompanied by remodeling of the central biosynthetic pathway of one carbon metabolism. The cells accumulated lipids while also inducing lipid uptake and synthesis genes, suggesting maladaptive metabolic rewiring.

**Conclusion:** Our findings indicate that glucose and lipids, the latter being the postnatally favored cardiac fuel, exert remarkably different consequences in MRPL44 deficient cardiomyocytes. The lipid enriched medium elicited robust activation of metabolic stress responses, with chronic upregulation of anabolic biosynthesis pathways and lipid accumulation indicative of conflicting metabolic homeostasis. These observations provide a mechanistic basis for postnatal disease manifestation and highlight nutrient metabolism as a key driver in development of infantile-onset mitochondrial hypertrophic cardiomyopathy.

**Highlights:**

- MRPL44 deficiency impairs mitochondrial translation but induces mtDNA replication and transcription in iPSC-derived cardiomyocytes.
- In glucose conditions, MRPL44 mutant cardiomyocytes upregulate mitochondrial replication and transcription program, but not translation.
- Lipid-enriched nutrient conditions exacerbate disease phenotype, inducing mitochondrial and ER stress responses in MRPL44 deficiency.
- Mitochondrial ribosome defect disrupts lipid homeostasis in cardiomyocytes causing impaired fatty acid oxidation, lipid accumulation and altered cholesterol metabolism.

## Introduction

Hypertrophic cardiomyopathy (HCM) is the second largest group of cardiometabolic disorders affecting more than 1.5 per 100000 newborn children. These inherited cardiomyopathies are characterized by impaired diastolic function and hypertrophic thickening of left ventricle, causing contractile defect, heart failure and sudden cardiac death [1,2]. Studies of infantile cardiomyopathy cohorts have identified pathogenic variants affecting mitochondrial (mt)translation as an important cause of neonatal and infantile HCM [3,4]. Mutations in mt-ribosomal subunit (*MRPL3, MRPL44*), tRNA processing, modification and aminoacylation (*ELAC2, GTPBP3, MTO1, TRMT5, AARS2*) and mitochondrial translation elongation factors (*TSFM, TUFM*) have all been linked to mitochondrial HCM (mHCM) [5–13]. The direct downstream effect of impaired mt-translation is oxidative phosphorylation (OXPHOS) deficiency, as mtDNA encodes 13 core subunits of OXPHOS complexes. Therefore, mt-translation defects can compromise both oxygen utilization and ATP production. However, primary defects affecting structural OXPHOS subunits, directly affecting the ATP synthesis machinery most often affect the brain rather than the heart [14,15]. The clinical and genetic evidence suggests that OXPHOS deficiency alone cannot fully explain mHCM, but the pathology may also involve altered metabolic signaling in cardiac development [15].

During the perinatal and neonatal period, the heart undergoes extensive metabolic maturation driven by developmental shift in substrate utilization [16,17]. Glucose and lactate are the primary fuel sources during the fetal development when the tissues experience low oxygen availability. Postnatally with onset of normoxic conditions, a rapid transition occurs towards mitochondrial fatty acid oxidation [16,17]. This switch to energy-efficient fatty acid metabolism is crucial for meeting the increasing energy demands of the maturing heart [16,17]. This metabolic transition in mammalian heart occurs during the early weeks after birth and is accompanied by increase in mitochondrial biogenesis, structural maturation of mt-proteome and crista remodeling, processes that are dependent on a functional mt-translation system [17,18]. Defects in this system disrupts the energy balance and hinders the metabolic adaptation to independent postnatal life.

In this study, we investigated the consequences of a mt-ribosome defect caused by a homozygous pathogenic mutation in MRPL44, a protein component of the large ribosomal subunit of the mt-ribosome [6]. Using MRPL44-mutant and control human cardiomyocytes differentiated from induced pluripotent stem cells (iPSCs), we examined pathogenic processes in fetal fuel conditions (glucose-medium) and postnatal ones (lipid-enriched medium). We show that patient-derived cardiomyocytes maintain homeostasis when glucose is available. However, in lipid-rich media designed to mimic postnatal conditions, the cells activate stress responses, including major alterations in cellular growth and metabolic pathways. These findings provide important insights into how mt-translation defects impair cardiomyocyte maturation and function, particularly during the critical transition from prenatal to postnatal life.

## Methods

### Ethical aspects

Skin biopsy samples were obtained from the patient and healthy volunteers after informed consent of the parents, following principles of Helsinki Declaration. The study work was approved by Helsinki and Uusimaa Hospital District ethical board.

### iPSC reprogramming

The fibroblasts were cultured from skin biopsy samples using standard protocol. iPSCs were generated by electroporating the fibroblasts with episomal plasmids. Briefly, the pCXLE-cDNA vectors (pCXLE-hOCT3/4-shp53-F, pCXLE-hSK and pCXLE-hMLN) were electroporated into dermal fibroblasts at 1650 V, 10 ms and 3 pulses with the Neon electroporator. Fresh fibroblast medium consisting of DMEM high glucose (#BE12-614F/12, Lonza), 10 % FBS (#F9665, Sigma-Aldrich) and L-Glutamax (#35050038, Thermo Fisher Scientific) was added on day 2 and day 4. On day 6, electroporated cells were replated on Matrigel (#354277, Corning) -coated plates containing fresh fibroblast media. On day 7, culture media was changed to Essential 8 media (#A1517001, Life Technologies) and was changed to fresh media every second day. Cells were maintained on E8 media, and the iPSC colonies were passaged mechanically by cutting on day 20-30.

### Differentiation of iPSCs to CMs and purification

iPSCs were differentiated to cardiomyocytes as in Burridge et al. [19] with the following modifications. iPSCs (>passage 20) were split at 1:10 or 1:12 ratios, using 0.5 mM EDTA (#15575020, Invitrogen) and grown for 4 days on growth factor reduced Matrigel (#356230, Corning), at which time they reached ∼85% confluence. Growth medium was changed to CDM3 media, consisting of RPMI 1640 medium (11875, Life Technologies), 500 µg/ml *O. sativa* –derived recombinant human albumin (#A0237, Sigma-Aldrich) (75 mg/ml stock solution was prepared in water for injection (WFI) quality H2O and stored at −20 °C), 213 µg/ml L-ascorbic acid-2-phosphate (Sigma-Aldrich, 64 mg/ml stock solution in WFI H2O, stored at −20 °C), 1 mM pyruvate (#11360039, Gibco) and 25 mg/ml uridine (#6680, Calbiochem). For day 0 to day 2, medium was supplemented with 3-6 µM CHIR99021 (#C-6556, LC Laboratories). On day 2, medium was changed to CDM3 supplemented with 2 µM Wnt-C59 (#S7037, Selleck Chemicals). Medium was changed to CDM3 media on day 4 and every other day (48hr). Contracting cells were observed from day 8 onwards.

iPSC derived CMs were purified using magnetic beads as per manufacturer’s instructions (#130-110-188, Miltenyi Biotec) on day 15-20 of differentiation. Cells were replated on growth factor reduced matrigel after purification and maintained in CDM3 media before treatment with different media conditions.

### Glucose or fatty acid medium for iPSC-CMs

Cells treated with normal glucose (GM) were changed to media with RPMI 1640 medium without glucose (#11879-020, Gibco) and 2 g/L glucose (#A2494001, Gibco) was added along with other constituents of CDM3 media for 72 hrs on day 27 of differentiation.

The CDM3 media was changed to fatty acid media (FM) on day 27 of differentiation for 72 hrs consisting of RPMI 1640 medium without glucose (#11879-020, Gibco), 1 g/L glucose (#A2494001, Gibco), 100 µM oleic acid (#03008-5ml, Sigma-Aldrich), 50 µM palmitic Acid (#P0500-10G, Sigma-Aldrich) along with other constituents of CDM3 medium.

### Immunofluorescence analysis

iPSCs or iPSC-derived CMs were plated on coverslips and allowed to grow for 3 days. Cells were fixed with 4% PFA for 15 minutes at room temperature, permeabilized with 0.1% Triton-X (Sigma-Aldrich), blocked with 10 % horse serum/PBS for 45 minutes at room temperature, and stained with primary antibodies: OCT4 (1:500 sc-8628, Santa Cruz), TRA-1-60 (1:500 MA1-023, Thermo Fisher), SSEA-4 (1:1000 MAB4304, Millipore ), SOX17 (1:500 AF1924, R&D systems), α-smooth muscle actin (1:500 A2547, Sigma), β-tubulin III (1:500 Ab18207, Abcam), 1:500 TNNT2 (ab45932, Abcam), TOM20 (#sc-17764, Santa Cruz Biotech) overnight at 4°C in 1% Horse serum /0.1 % TritonX /1 % BSA/PBS. Cells were washed three times for 5 minutes in 1 % BSA/PBS and then incubated for 1 hr at room temperature in the dark with secondary antibodies 1:400 Alexa Fluor 488 (A-21206, Invitrogen), 1:400 Alexa Fluor 594 (A11005, Invitrogen) in 1% BSA/PBS. In addition, for neutral lipid droplet staining, the samples were incubated with Lipidtox red (H34476, Invitrogen) for 30 minutes. Cells were washed again with PBS and mounted with Vectashield mounting medium with DAPI, to stain the nuclei (H-1200, VectorLabs). Images for OCT4, TRA-1-60, SSEA-4, SOX17, α-smooth muscle actin and β-tubulin III were acquired by EVOS FL cell imaging system (Thermo Fisher). Images for TNNT2, TOM20 and Lipidtox were acquired by fluorescence confocal microscope (LSM880) with 40x objective.

### Image quantification

Mitochondrial area and cell area were quantified using ImageJ [20]. Cell area was determined from TNNT2 immunostaining. After applying an intensity threshold, cell boundaries were outlined, and the resulting regions of interest (ROIs) were quantified using the “Measure” tool in ImageJ. These ROIs were then applied to the mitochondrial channel, where mitochondria were segmented by thresholding. The “Measure” tool was used to quantify the mitochondrial area within each ROI. Lipid droplet images were quantified using CellProfiler software [21]. Cell boundaries were identified from TNNT2 immunostaining, and lipid droplets were segmented from the Lipidtox channel using the “IdentifyPrimaryObjects” module. Parent-child relationships were established between cells (parents, defined as cell boundaries) and lipid droplets (children), enabling quantification of droplet number per cell.

### Karyotyping

Cells were split one or two days before collection. Hypotonic solution containing 0.0075 M KCl and 0.2-0.4 ml colchicine was added to arrest the cell cycle in metaphase. Methanol / glacial acetic acid was used in 3:1 ratio for fixation. The chromosomes were visualized after Giemsa staining and analyzed by light microscopy.

### FACS analysis

Cells were transferred to flow cytometry tubes (BD Biosciences) and fixed with 4% PFA for 15 min, permeabilized with 90% methanol for 15 min, and stained with 1:200 TNNT2 (ab45932, Abcam) for 45 min at room temperature. Secondary antibody was 1:500 Alexa Fluor 488 (A-21206, Invitrogen). At least 26000 cells were analyzed using Sony SH800S cell sorter with 100 μm microfluidic sorting chip.

### Quantitative RT-PCR

Total RNAs were purified using Trizol reagent (Invitrogen). Remnant genomic DNA was eliminated by DNAse treatment and cDNA was synthesized using the manufacturer’s instruction (#K1672, Thermo Fisher Scientific). Real-time quantitative PCR (RT-qPCR) reactions were performed using SensiFast No-Rox kit (#BIO-98020, Bioline) using CFX96 Touch qPCR system (Bio-Rad). Relative expression of a transcript was determined by normalizing to the expression of housekeeping genes beta-actin.

NCBI primer BLAST software [22] was used for oligonucleotide design, 70–150 bp-long product was amplified, and oligonucleotides were designed to be separated by at least one intron of the corresponding genomic DNA of a typical minimum of 1,000 bp length. Primers used in this study are mentioned in Table S3.

### mtDNA copy number analysis

A standard proteinase K and phenol-chloroform extraction method was used to isolate total cellular DNA from frozen cell pellets. mtDNA copy number was measured by quantitative PCR (qPCR) by amplification of mtDNA fragment MT-CYTB against nuclear-DNA-encoded APP gene. 25 ng of DNA was used per PCR reaction, all qPCR assays were run in triplicates on CFX96 Touch qPCR system (Bio-Rad).

### RNAseq analysis

RNA was extracted from purified cardiomyocytes preparations using RNA-binding columns (#74106, Qiagen RNeasy Kit) as per manufacturer’s instructions. RNA was analyzed using TapeStation, and samples with RNA integrity number over 7 were used. RNA samples were sequenced in Biomedicum Functional Genomics Unit at the Helsinki Institute of Life Science and Biocenter Finland (FUGU), University of Helsinki. The “Bulkseq” 3’ UTR RNA sequencing was conducted on iPSC-CMs samples, as described in [23]. Differential expression analysis was done using the DESeq2 package in R environment with default settings. The count values were normalized between samples using a geometric mean. Sample-wise factors were estimated to correct for library size variability and estimation of dispersion (i.e. variance, scatter) of gene-wise values between the conditions. A negative binomial linear model and Wald test were used to produce p-values. Low-expression outliers were removed using Cook’s distance to optimize the p-value estimation and finally, multiple testing adjustment of p-values was done with the Benjamini-Hochberg procedure.

### Bioinformatics analysis

Pathway analysis was performed for significantly upregulated genes (p-value <0.05), or, separately significantly downregulated genes (p-value <0.05) using Qiagen Ingenuity Pathway analysis [24] using default settings. The top 10 pathways were selected for representation in each case. Comparison analysis from Ingenuity Pathway Analysis (Qiagen) was used to predict pathway changes in different treatment conditions.

Cis-regulatory analysis was performed with iRegulon [25] (v. 1.3) in Cytoscape (v.3.7). The input was most significantly upregulated genes (p<0.01), or separately significantly downregulated genes (p<0.01). For enrichment of transcription factor binding motifs, the human gene list was used as-is. In all cases, the regulatory regions interrogated comprised 10 kbp upstream and 10 kbp downstream of the transcription start site, as annotated in iRegulon. The seven species option was used in the rankings of transcription factor targets. The ROC threshold for AUC calculation was set at 1%.

### Statistical analysis and data visualization

Statistical analyses were performed using Prism 8 software and R; graphs were made with Prism 8 and R. padj and p-values were used throughout the study to determine statistical significance of these comparisons. Volcano plots and Box plots were done using “EnhancedVolcano” and “ggplot2” function in R.

## Results

### Generation of MRPL44 iPSCs and differentiation to cardiomyocytes

Cardiomyocytes from induced pluripotent stem cells (iPSCs) were generated from patient fibroblasts carrying homozygous pathological variant in *MRPL44* gene (c.467T>G, p.L156R) and from three healthy voluntary controls (C1, C2 and C3) (study approach, Figure. 1A). The presence of the mutation in the patient cells was confirmed by sequencing the corresponding coding region (Figure. S1A). Quantitative RT-PCR and immunostaining demonstrated the expression of pluripotent marker genes in all generated iPSC lines (Figure. 1B, S1B and S1C) and silencing of the episomal vectors (Figure. S1D). All the cell lines showed normal karyotypes (Figure. S1E). All the cell lines were able to differentiate into three germ layers indicating their pluripotency (Figure. S1F). These findings indicate successful generation of iPSC lines from both patients and control-derived fibroblasts.

**Figure 1.**
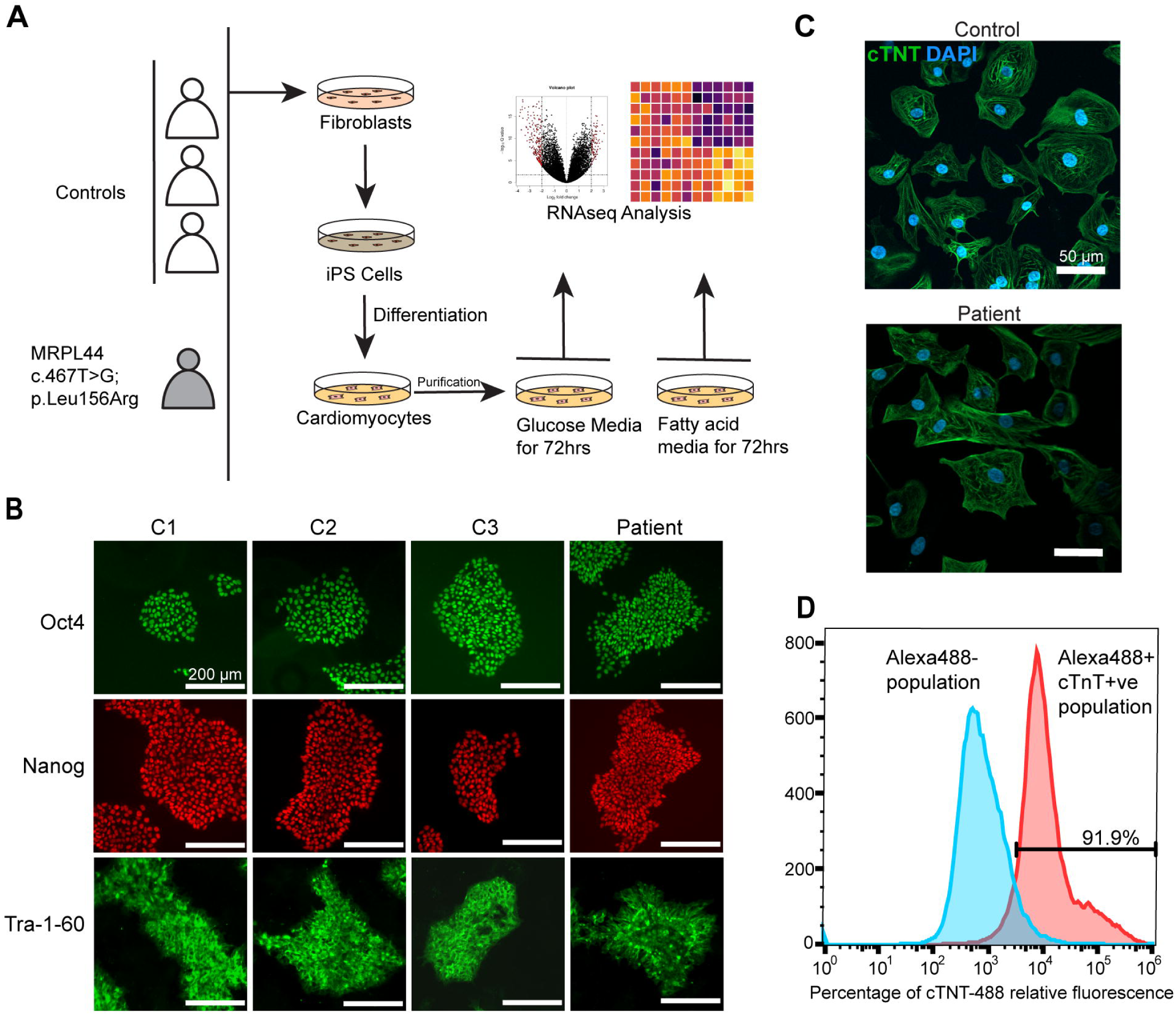
**Generation and characterization of iPSC-derived cardiomyocytes.** A. Schematic overview of the experimental design. B. Reprogramming factor expression in control and MRPL44 patient-derived iPSCs after reprogramming (OCT4; green, NANOG; red and TRA-1-60; green) (Scale bar; 200µm) C. Cardiac marker positivity of iPSC derived CMs. Immunofluorescence, cardiac troponin (cTnT; green) and nuclei (DAPI, blue) (Scale bar; 50 µm). D. Purification of iPSC derived cardiomyocytes. Representative FACS analysis shows 91.9% cTnT positive cells after purification.

We further differentiated the iPSCs into cardiomyocytes as a monolayer culture using a small molecule-based induction of differentiation protocol as described previously [19] (Figure. 1A). After 8-10 days of differentiation, all the iPSC-derived cardiomyocyte lines (from now on, CMs) exhibited widespread spontaneous beating activity (Supplementary Movies 1-4). Immunofluorescence and FACS analysis indicated that >80-90 % of cells were positive for cardiac troponin T (Figure. 1C and D), indicating successful cardiomyocyte differentiation.

### Mitochondrial translation defect drives transcriptional reprogramming of genes encoding mitochondrial proteins in iPSC-derived cardiomyocytes

To elucidate the pathogenesis-related changes in patient CMs, we performed RNA sequencing on patient and control -derived CMs cultured in glucose-rich medium (GM, 2 g/L). The most upregulated transcript was *LINC00551*, a ferroptosis regulator, and most downregulated mitoregulin (*MTLN*), a mitochondrial protein involved in very long-chain fatty acid oxidation (Figure. 2A). The overall expression of transcripts encoding mitochondrial proteins were markedly elevated, indicating mitochondrial biogenesis induction (Figure. 2A, B and Table S1). The expression of the mtDNA-encoded transcripts, particularly those encoding subunits of OXPHOS complexes I and V, showed the strongest induction (Figure. 2A–C). The mRNAs of nuclear OXPHOS subunit genes were modestly increased (Figure. 2C). The patient CMs showed an increase of 2.5-fold in mtDNA copy number compared to controls (Figure. 2D), and mitochondrial content per cell was also increased (Figure. 2E). Despite this transcriptional upregulation, steady-state complex I protein level was markedly reduced (Figure. 2F), indicating that MRPL44 deficiency activated replication and transcription of mtDNA, which could not, however, compensate for the translation defect in CMs.

**Figure 2.**
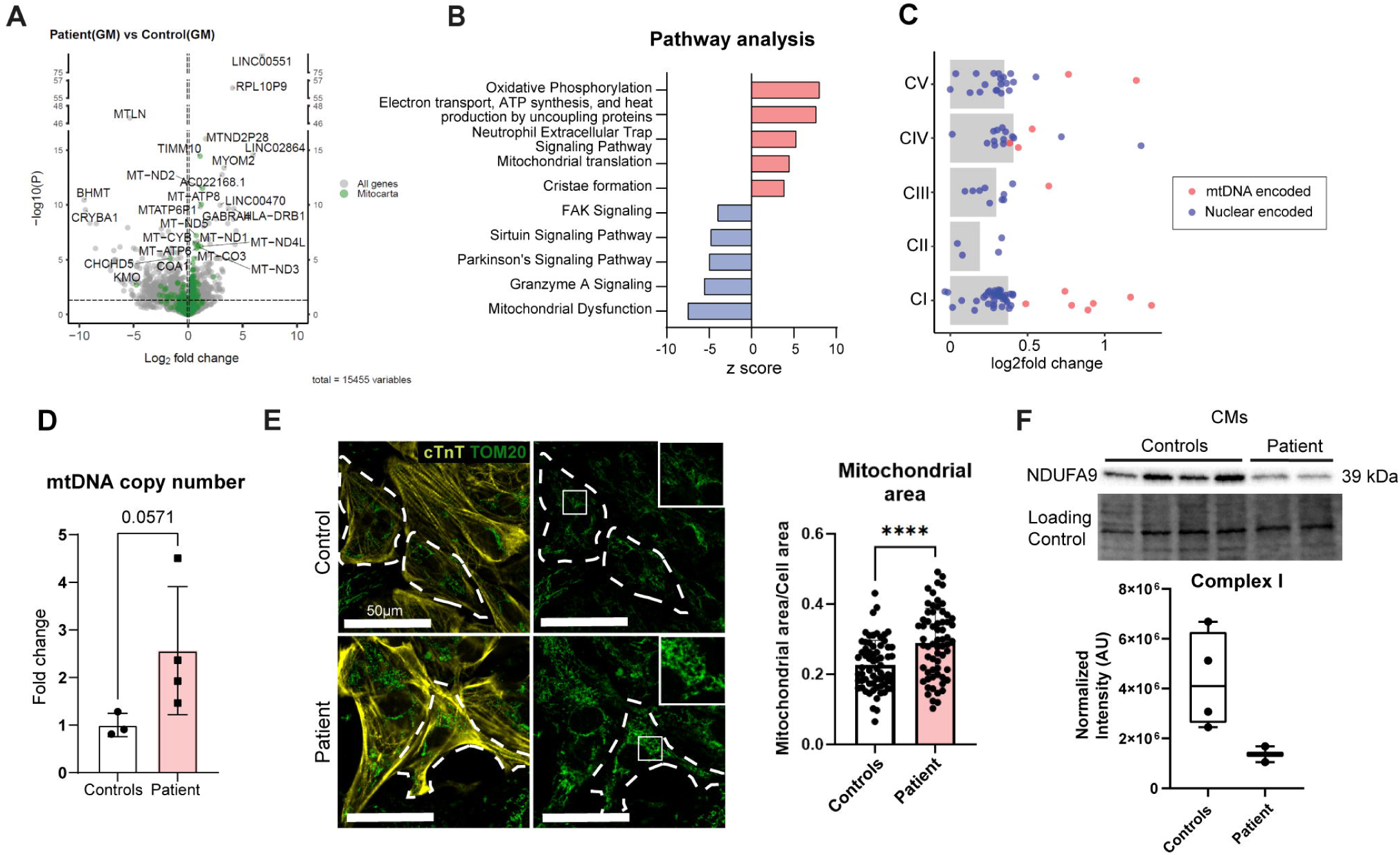
**Mitochondrial transcriptome in patient CMs under glucose medium (GM) conditions.** A. Mitochondrial transcriptome. Volcano plot showing the fold changes and statistical significance of the transcriptomic changes in patient CMs in GM conditions compared to control. Green dots: genes encoding MitoCarta (mammalian mitochondrial protein database) proteins. Dotted line: significance threshold at P<0.05 (Wald test); n ≥ 5 independent differentiation experiments for three control cell lines and one patient cell line. B. Top affected pathways identified by Ingenuity Pathway Analysis program from significantly changed genes of patient CMs vs controls (P<0.05) (Fisher’s exact test). C. OXPHOS gene expression levels. Bar plot: fold changes in patient CMs. Blue dots: nuclear encoded genes; red dots, mtDNA encoded genes. D. mtDNA copy number in control and patient CMs. Independent differentiation experiments of controls and patient CMs, n ≥3. Student’s t-test. E. Mitochondrial aggregation in patient CMs. Representative image. (Cardiac troponin, cTnT, yellow; TOM20, green; DAPI, blue); Scale bar; 25 μm. Right-side panel: increased mitochondrial area per cell in patient CMs. Student’s t-test. F. Mitochondrial complex I (NDUFA9) protein amount. Western Blot analysis of patient and control cardiomyocytes.

To mimic the perinatal metabolic switch in cardiac tissue from fetal glucose-dependent metabolism to postnatal lipid-based metabolism we cultured CMs in low-glucose medium (1 g/L) and supplemented it with two major fatty acids, palmitic acid and oleic acid, the lipid species prevalent in the newborn heart [26]. In the fatty acid enriched medium (FM), control CMs showed activation of lipid oxidation and uptake (Figure. 3A) along with induction of OXPHOS components (Figure. 3F), since lipids enhance mitochondrial oxidative metabolism during cardiac maturation. In addition, controls showed higher expression of cardiomyocyte maturation markers, including metabolic (*CKMT2, PPARGC1A, PDK4*), electrophysiological (*KCNJ2*) markers and lower expression of immature cardiomyocytes marker (*MYH6*) (Figure. 3A and S2A).

**Figure 3.**
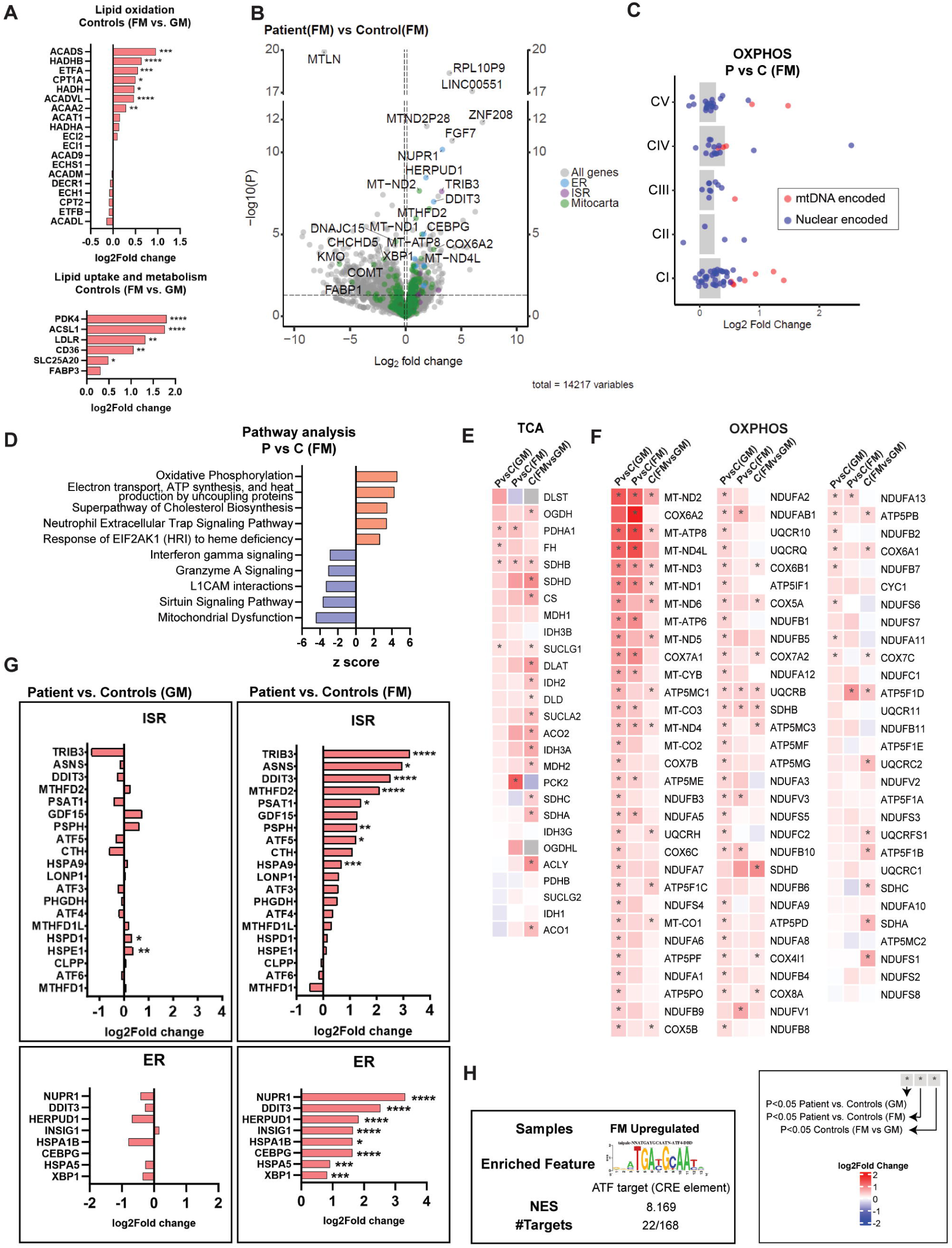
**Induction of stress response pathways in patient CMs in fatty acid media (FM) conditions** A. Expression of lipid oxidation and uptake genes in control CMs in fatty acid enriched medium. B. Volcano plot showing the fold changes and statistical significance of the transcriptomic changes in patient CMs in FM conditions. Mitocarta gene expression, green; ER stress, blue; ISRmt, purple dots. P<0.05; Wald test; n ≥ 3 independent differentiation experiments for three control cell lines and one patient cell line C. Expression of OXPHOS complex genes, FM conditions. Bar plot: fold changes of OXPHOS complex gene expression in patient CMs. Blue dots: nuclear encoded genes; red dots: mtDNA encoded genes. D. Top affected pathways identified by Ingenuity pathway analysis from significantly changed transcripts of patient CMs in FM conditions. P<0.05; Fisher’s exact test. E. Heatmap of transcripts encoding TCA cycle components; three-way comparisons (patient vs control CMs in glucose conditions; patient vs control CMs in fatty acid conditions; control CMs in FM vs. GM conditions. *P ≤ 0.05; Wald test. F. Heatmap of transcripts encoding OXPHOS components in three-way comparisons as in E. Significantly changed transcripts marked with asterisk; *P ≤ 0.05, Wald test. G. ISRmt and ER stress response gene expression. H. cis-regulatory motif analysis. Enriched binding motifs in the promoters of the significantly upregulated genes of patient CMs in FM conditions (P<0.01). iRegulon analysis of transcription factor binding motifs and ChIP-seq peaks; NES, normalized enrichment score; # Targets, number of gene targets.

The patient cardiomyocytes cultured in FM exhibited pronounced induction of transcripts of mtDNA-encoded OXPHOS subunits (Figure. 3B–D, F and Table S1). OXPHOS assembly factors exhibited a split response: genes for cofactor handling (*COX19* -copper, *NUBPL* – iron-sulfur clusters, *COX15* - heme, *COA7* – heme binding, *COX10* - heme) showed a trend of downregulation or no significant change while others participating in assembly or import of protein complex subunits were typically increased in expression (Figure. S2B). Components of mitochondrial translocases and chaperones for mitochondrial protein import (*TIMM8A, TIMM21, TIMM10*) and sorting were upregulated in both conditions and proteins for cristae organization were also increased (Figure. S2C, D).

Pyruvate dehydrogenase subunit A *(PDHA1),* a key enzyme of oxidative glucose usage enabling pyruvate entry to tricarboxylic acid (TCA) cycle, was induced under both nutrient conditions, indicating the dependence on glucose oxidation even in fatty-acid enriched, glucose depleted medium. TCA cycle enzymes were induced with an upward trend in patient and control cardiomyocytes under both GM and FM conditions. Cis-regulatory analysis of significantly upregulated genes, using publicly available ChIP-seq datasets [25], identified estrogen related receptor alpha (*ESRRA*) transcription factor as an upstream regulator with an enriched binding motif for the upregulated genes in patient CMs (Figure. S2E, Table S2). ESRRA has been shown to be a transcriptional coactivator for genes involved in OXPHOS and mitochondrial biogenesis [27].

Taken together, these findings indicate that the defect of mitochondrial ribosome subunit MRPL44 in cardiomyocytes triggers a coordinated transcriptional reprogramming in cardiomyocytes promoting sustained glucose usage.

### Fatty acid conditions trigger mitochondrial and ER stress responses in patient CMs

The most prominent change in patient CMs under FM was activation of multiple components of the mitochondrial integrated stress response (ISRmt), a key metabolic and transcriptional program triggered by mitochondrial dysfunction, particularly mtDNA expression defects. The response is regulated by ATF transcription factor family, whose targets were the top hits in the motif enrichment analysis (Figure. 3H, S2F, S2G and Table S2). In patient CMs, *ATF5* was increased and along with its targets, including ISRmt related genes tribbles pseudokinase 3 (*TRIB3*), asparagine synthetase (*ASNS*), mitochondrial methylene tetrahydrofolate dehydrogenase (*MTHFD2*), and metabokine *GDF*15 (growth differentiation factor 15.) Furthermore, ISRmt-linked transcription factor DNA-damage-inducible transcript 3 (*DDIT3*), de novo serine biosynthesis genes (*PSAT1*, *PSPH*), CCAAT enhancer binding protein gamma (*CEBPG*) were significantly increased in patient CMs (Figure. 3G). However, out of the typical signature of ISRmt genes, *FGF21* response was absent (Figure. S2H).

Parallel to ISRmt activation, markers of ER stress were also notably elevated in FM conditions (Figure. 3G). These included *HERPUD1*, *NUPR1*, *SREBP1*, *INSIG1*, *XBP1*, *CHAC1*, and *HSPA5*.

Although *XBP1* expression increased, stress-induced splicing was absent, indicating partial ER stress activation (Figure. S2I). Genes linked with ferroptosis and/or antioxidant defense systems were increased in expression. Especially in FM, glutathione peroxidase 4 (*GPX4*), glutathione peroxidase 7 (*GPX7*) and metallothionein 1X (*MT1X*) as well as nuclear protein 1, transcriptional regulator (*NUPR1*) were upregulated. Additionally, Fibroblast growth factor 7 (FGF7), related to mitigating oxidative stress in myocardial injury was induced in patient CMs in FM conditions (Figure. 3B).

In glucose medium, patient cardiomyocytes showed a trend toward upregulation of mitochondrial ribosomal genes, but these changes were absent when fatty acids were the fuel (Figure. S3A). While cytosolic ribosomal genes were not induced in FM, several cytoplasmic tRNA synthetases (*TARS1*, *IARS1*, *AARS1*, *SARS1*, *YARS*) were upregulated (Figure. S3C), consistent with amino acid deprivation. Mitochondrial tRNA synthetases were unchanged (Figure. S3B). Ingenuity Pathway Analysis predicted nutrient-dependent alterations in mTOR signaling and glucose metabolism in FM compared to GM (Figure. S3D). *NMRK2*, a key enzyme in NAD[ salvage, and *SLC25A11*, an oxoglutarate/malate carrier mediating mitochondrial NADH shuttling, were both induced (Figure. S3E), suggesting altered NAD metabolism.

These results demonstrate that the translational defect due to MRPL44 mutation prevents cardiomyocytes from effectively transitioning to fatty acid metabolism. This metabolic stress induces mitochondrial and ER stress pathways with signs of amino acid imbalance.

### Lipid homeostasis is affected in patient CMs

The pathology associated with lipid-enriched nutrition and previous autopsy data of cardiac lipid accumulation in the MRPL44 patient [6], prompted us to directly examine lipid droplets in CMs. Neutral lipid staining revealed a marked increase in lipid droplet content in patient CMs compared to controls (Figure. 4A and B). Transcriptomic profiling revealed aberrant lipid uptake and metabolic signatures: induction of *CD36* (lipid transporter), *LDLR* (cholesterol endocytosis), *ACSL1* (long-chain fatty acid-CoA ligase), *PDK4* (inhibitor of glycolytic pyruvate oxidation), *ASAH1* (acid ceramidase) (Figure. 4C and S4A, B) and *MTLN*, a mitochondrial outer membrane protein involved in catabolism of very long-chain fatty acids (Figure. 2A and 3B). Especially in FM, a robust upregulation of cholesterol biosynthesis genes, including *HMGCR*, *HMGCS1*, *FDFT1*, *SQLE*, and acetyl-CoA acetyltransferases (*ACAT1*, *ACAT2*) was found in MRPL44 deficient cells (Figure. S4C). Control CMs activated this pathway to a lesser extent, and some cholesterol biosynthesis genes, such as *SC5D* and *NSDHL*, were also elevated in patient CMs fed with glucose (Figure. S4C).

**Figure 4.**
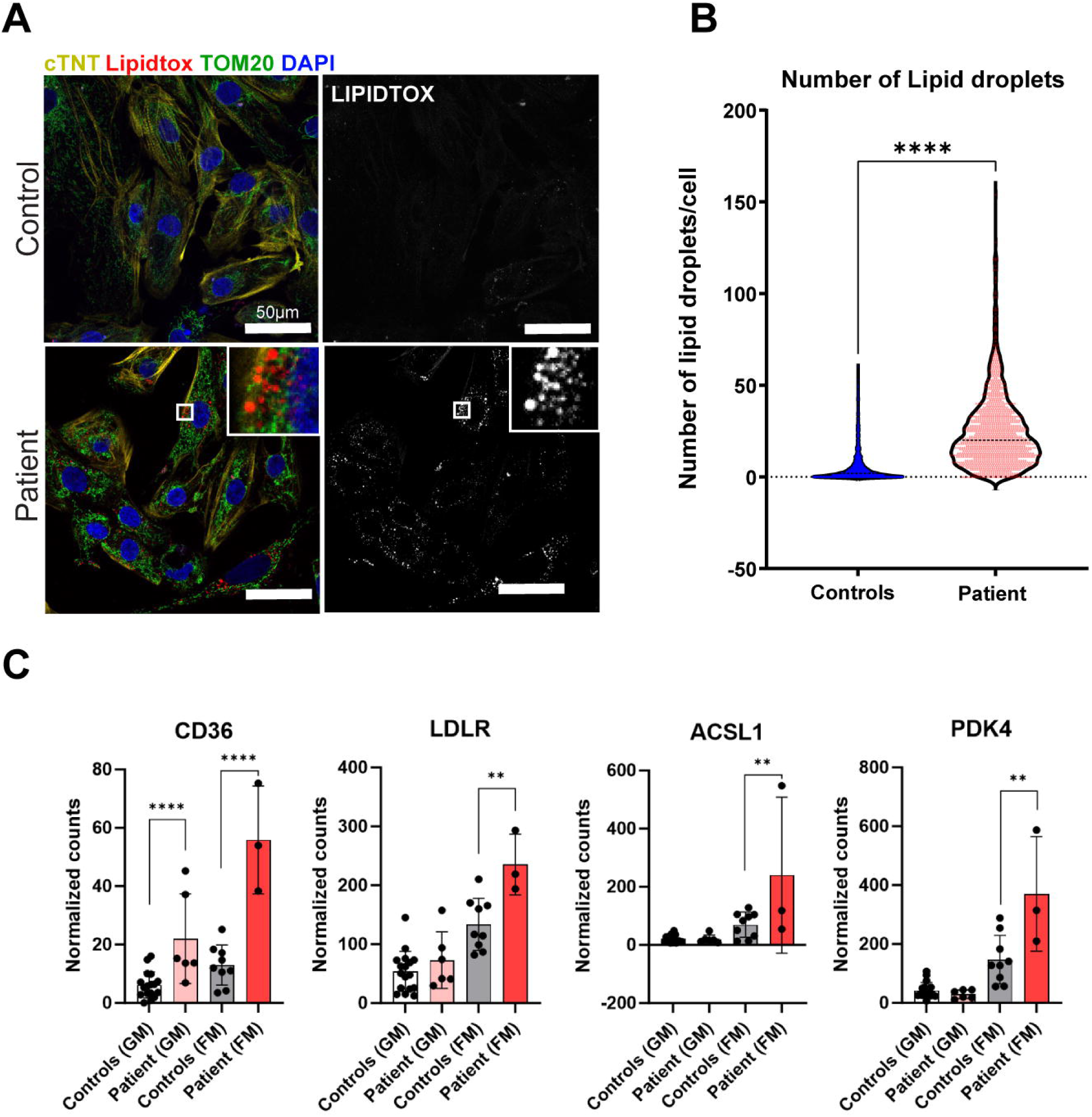
**Impaired lipid homeostasis in patient CMs** A. Lipid droplet analysis. Immunostaining of neutral lipid droplets in MRPL44 and Control iPSC-CMs by Lipidtox. (cTnT, yellow; Lipidtox, red; TOM20, green; DAPI, blue; scale: 50 µm) B. Quantification of droplets per cell between patient and control CMs. (n=1273 CMs) *P ≤ 0.05, **P ≤ 0.01, ***P ≤ 0.001, ****P ≤ 0.0001 (Student’s t-test) C. Gene expression levels of lipid transporter *CD36, LDLR, ACSL1* and *PDK4* gene in different media conditions. *P ≤ 0.05, **P ≤ 0.01, ***P ≤ 0.001, ****P ≤ 0.0001 (Wald test)

Together, these findings indicate a profound disruption of lipid homeostasis in MRPL44-deficient CMs, characterized by reduced lipid utilization, increased lipid uptake, imbalanced ceramide and alteration in cholesterol metabolism with capacity to exacerbate cardiomyocyte dysfunction.

## Discussion

Mitochondrial translation defects are an important cause of hypertrophic cardiomyopathy in infants, although the underlying patho-mechanism remains poorly understood [28,29]. Mitochondrial large ribosomal subunit 44 protein deficiency is one of the mitochondrial causes of HCM. Here, we report the molecular consequences of *MRPL44* defect in iPSC-derived cardiomyocytes of a patient with progressive mitochondrial HCM with especial interest in nutrient effects. We show that maladaptive metabolic responses are particularly evident when the cells are cultured in fatty acid enriched media, the favored nutrient in healthy post-natal heart [30]. The MRPL44 deficiency linked metabolic remodeling included increased lipid uptake, altered cholesterol biosynthesis and lipid droplet accumulation, together with downregulation of very long-chain fatty acid catabolism. The accumulation of lipids together with oxidative stress signatures are likely inducers of mitochondrial and ER stress. These, when chronically upregulated, lead to remodeling of metabolome, with potential progression to ferroptosis [31]. Importantly, the nutrient-dependent nature of these phenotypes provides a mechanistic explanation for the absence of overt prenatal pathology, as the fetal heart primarily relies on glycolytic glucose oxidation. However, inability to shift to lipid usage after birth is a potent driver of pathology with dramatic disease manifestation and progression during the first months of postnatal life.

The fact that a mitochondrial ribosomal defect, which compromises OXPHOS activity and energy metabolism, leads to an overgrowth of the cardiac tissue seems counterintuitive. However, the deficiency activates a persistent anabolic mitochondrial integrated stress response (ISRmt) in cardiomyocytes previously reported in mitochondrial muscle disease also to involve mTORC1, the master regulator of growth [32–38]. While inactive under glucose-rich conditions, this pathway became chronically activated upon lipid exposure, suggesting that sustained anabolic stress signaling during the perinatal metabolic shift drives pathological growth via mTORC1. ISRmt together with the amino acid tRNA synthetases support such activation. Previous studies have shown that in the mouse heart, deficiency of COX10, a component of cytochrome c oxidase, drives mTOR hyperactivation and ferroptosis. [39]. Together, the data suggest that perinatal ISRmt activation is maladaptive, promoting growth and hypertrophy.

We found that MRPL44 deficient cardiomyocytes activated mtDNA synthesis and mt-transcription, which did not, however lead to increase of OXPHOS complex I protein amount, with the largest number of mtDNA-encoded subunits and iron-sulfur clusters. Our observation that especially metal-linked assembly factors did not follow the general upregulation of other mitochondrial targeted transcripts could imply a regulatory feed-back loop to metal-inserting protein factors, to ensure their insertion to OXPHOS subunits only when mitochondrial ribosome is functional. While MRPL44 defect and different mitochondrial translation-linked cardiomyopathies are often deadly in early life, the surviving patients may stabilize by the age of 5-6 years [3,6]. How the lipid metabolic abnormalities and stress responses can be compensated by the growing heart, remains unknown but represents an important area for therapeutic exploration for these devastating disorders. Our findings suggests that lipid accumulation, increased lipid uptake and induction of metabolic stress responses are a central pathogenic mechanism of MRPL44-linked cardiomyopathy and present a metabolic cause for cardiac hypertrophy during the first years of life.

## Data availability

The RNA-seq datasets have been deposited to European nucleotide archive (ENA) with accession no. PRJEB74932.

## Contributions of authors

Conceptualization, S.P., N.A.K. and A.S.; Experimentation S.P., T.M., and N.A.K.; Data curation and analysis S.P., T.M, N.A.K., A.Z.; Interpretation, S.P., N.A.K., A.S., A.Z.; Manuscript writing and editing, all authors.; Supervision, A.S.; Funding Acquisition, S.P. and A.S. All authors have read and approved the final version.

## Acknowledgements

The authors would like to thank Markus Innilä and Tuula Manninen for technical assistance and Biomedicum Functional genomics unit (FuGU) for bulk RNAseq data analysis. We are grateful for the financial support of Research Council of Finland (to A.S. and N.A.K. #316435, 355637), Finnish Foundation for Cardiovascular Research, Sigrid Jusélius Foundation, Jane and Aatos Erkko Foundation (to A.S.), Finnish cultural foundation, Instrumentarium Foundation and University of Helsinki Doctoral school of Integrative Life Sciences (to S.P.).

## Abbreviations

HCM: hypertrophic cardiomyopathy
MRPL44: mitochondrial ribosome protein large subunit 44
OXPHOS: oxidative phosphorylation
iPSCs: induced pluripotent stem cells
iPSC-CMs: iPSC-derived cardiomyocytes
GM: glucose Media
ESRRA: estrogen related receptor alpha
FM: fatty acid Media
UPRmt: mitochondrial unfolded protein response
ER stress: endoplasmic reticulum stress
ATF4: activating transcription factor 4
ATF5: activating transcription factor 5
mTORC1: mammalian target of rapamycin 1
mtDNA: mitochondrial DNA
TRIB3: tribbles pseudokinase 3
ASNS: asparagine synthetase
MTHFD2: mitochondrial folate metabolism gene
GDF15: growth differentiation factor 15
DDIT3: DNA-damage-inducible transcript 3
CEBPG: CCAAT enhancer binding protein gamma
GPX4: glutathione peroxidase 4
NUPR1: nuclear protein 1 transcriptional regulator
GPX7: glutathione peroxidase 7
NMRK2: nicotinamide riboside kinase 2
GPX1: glutathione peroxidase 1
MTLN: mitoregulin
CD36: cluster of differentiation 36
LDLR: low-density lipoprotein receptor
ASAH1: N-acylsphingosine amidohydrolase 1

---

**Supplementary Figure 1.**
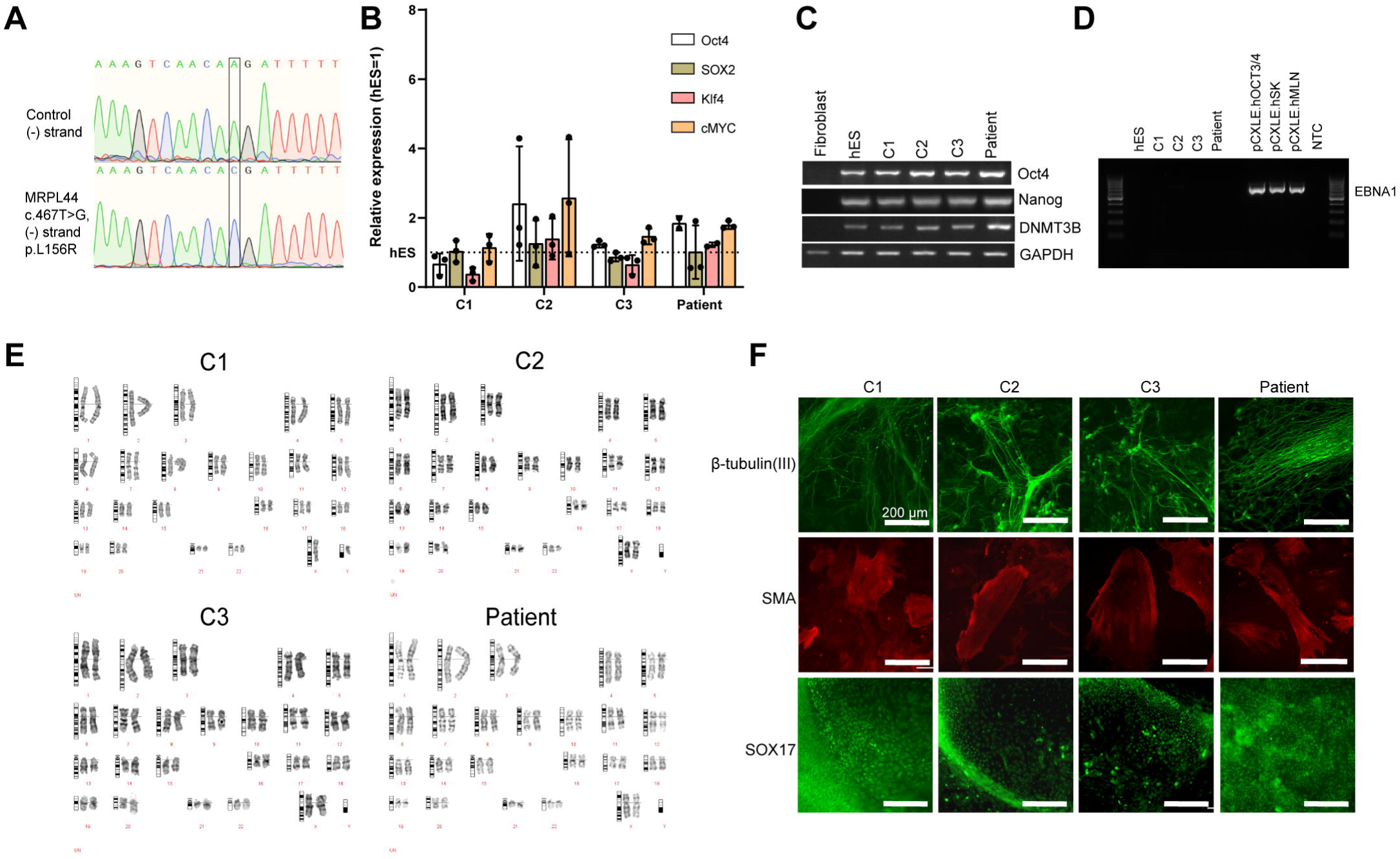
**Characterization of control and patient CMs** A. Homozygous mutation in patient CMs c.467T>G; p.L156R by Sanger sequencing B. Expression of reprogramming factors (*OCT4, SOX2, KLF4* and *c-MYC*) in control and patient iPSCs compared with human ES cell line. C. RT-PCR analysis of ES cell-specific transcripts (*OCT4*, *NANOG* and *DNMT3B*). D. Exogenous plasmid expression in the control and patient iPS cell lines by PCR with passage > 20 E. Karyotyping analysis of control cell lines and patient iPSCs F. Immunostaining images of embryoid bodies derived from control and patient iPSCs showing expression of β-III-tubulin (ectoderm; green), smooth muscle actin (SMA) (mesoderm, red) and Sox17 (endoderm, green). Scale bar; 200 μm.

**Supplementary Figure 2.**
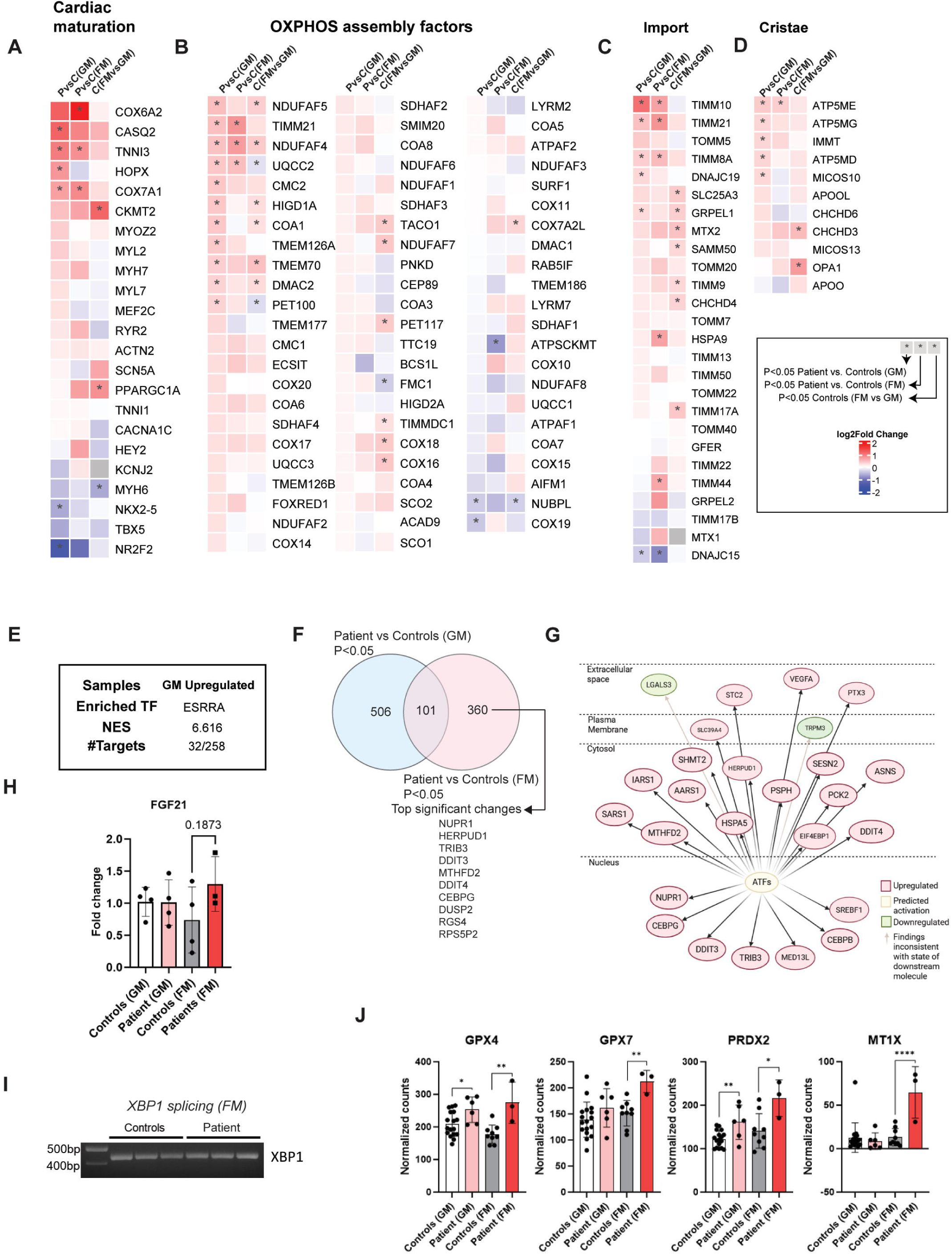
**FM media induces stress responses in patient CMs** A. Heatmap showing log fold changes in gene transcripts encoding cardiac maturation markers in three comparisons (Patient vs Control CMs in GM conditions, Patient vs Control CMs in FM conditions, Control CMs (FM vs. GM conditions)). Significant genes (P<0.05 (Wald test)) are marked with “*”. B. Heatmap showing log fold changes in gene transcripts encoding OXPHOS assembly factors. C. Heatmap showing log fold changes in gene transcripts encoding mitochondrial import genes. D. Heatmap showing log fold changes in gene transcripts encoding mitochondrial cristae genes. E. Cis-regulatory analysis. Enriched features in the promoters of the significantly changed genes of patient CMs in GM conditions (P<0.01), based on iRegulon analysis of transcription factor binding motifs and ChIP-seq peaks. TF, transcription factor; NES, normalized enrichment score; # Targets, number of gene targets F. Venn diagram showing commonly changed and differentially changed number of genes in GM and FM conditions in patient CMs and top 10 significant changes in patient CMs in FM (P<0.05) G. Schematic showing differentially changed genes known to be regulated by ATFs in FM conditions based on IPA analysis H. *FGF21* gene expression analyzed by q-RTPCR I. *XBP1* splicing of control and patient CMs in FM conditions. *XBP1* gets spliced upon prolonged endoplasmic reticulum stress, but is intact in patient CMs J. Expression of oxidative stress genes (*GPX4, GPX7, PRDX2* and *MT1X*)

**Supplementary Figure 3.**
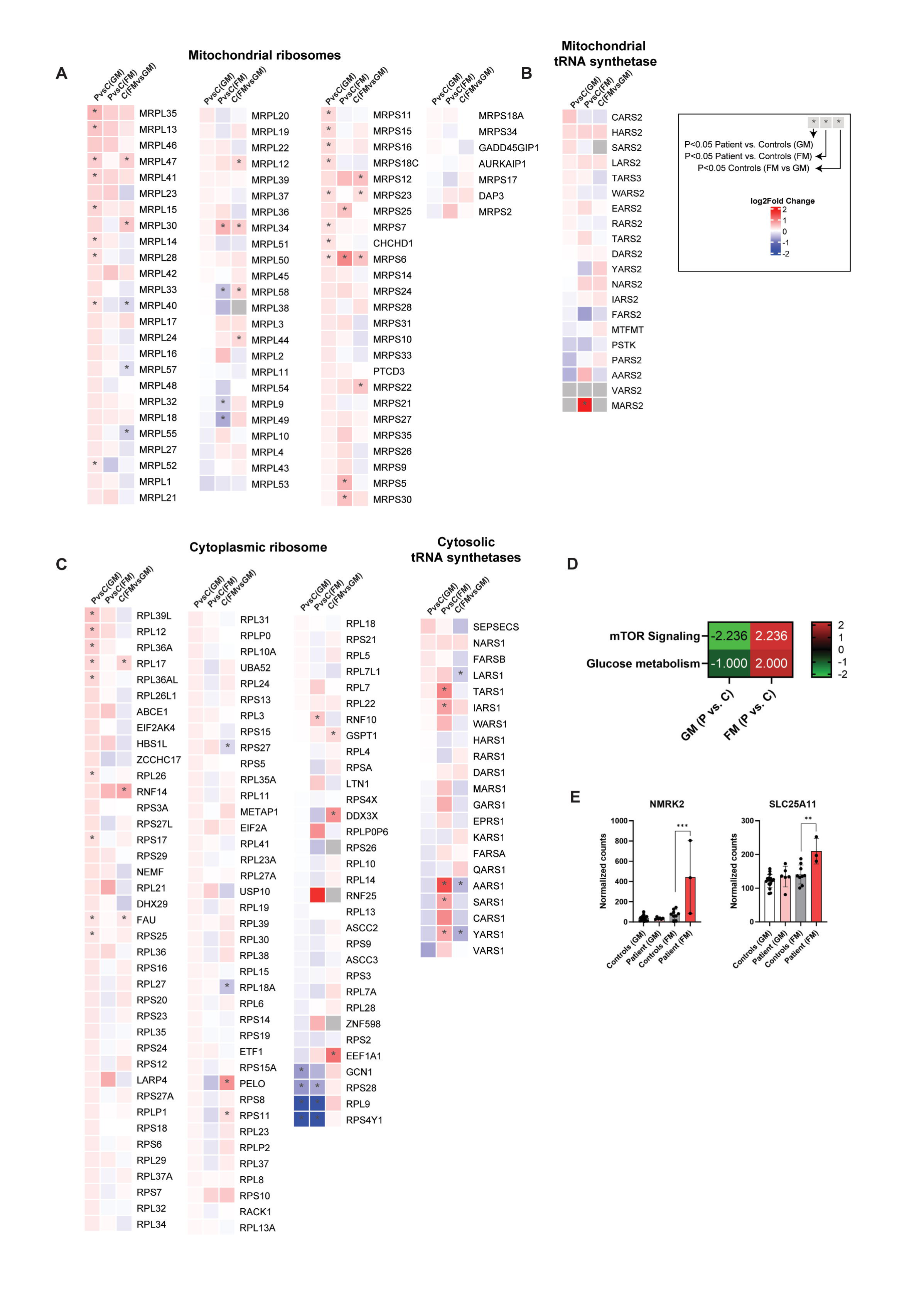
**Mitochondrial transcriptome and stress responses in FM media** A. Heatmap showing log fold changes in gene transcripts encoding mitochondrial ribosome components in three comparisons: patient vs control CMs in GM conditions; patient vs control CMs in FM conditions; control CMs (FM vs. GM conditions). Significant genes (P<0.05 (Wald Test)) are marked with “*”. B. Heatmap showing log fold changes in gene transcripts encoding mitochondrial tRNA synthetases. C. Heatmap showing log fold changes in gene transcripts encoding cytoplasmic ribosome components and cytoplasmic tRNA synthetases. D. Comparison analysis from Ingenuity pathway analysis shows predicted activation of mTOR signaling and glucose metabolism change in FM conditions. E. Gene expression of *NMRK2* and *SLC25A11* from RNAseq analysis.

**Supplementary Figure 4.**
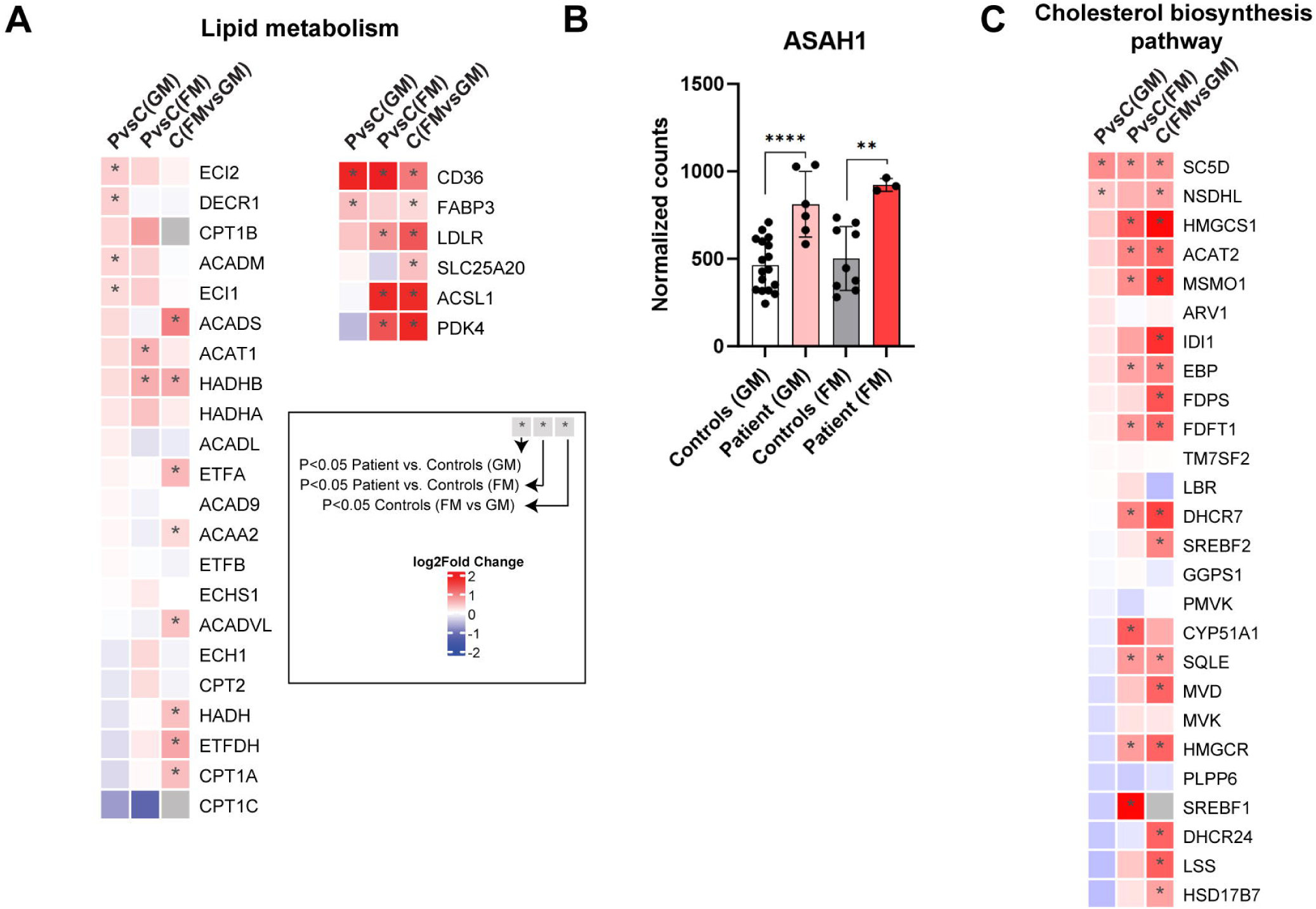
**Alteration of lipid metabolism in patient CMs** A. Heatmap showing log fold changes in gene transcripts encoding lipid oxidation and lipid uptake components in three comparisons: patient vs control CMs in GM conditions; patient vs control CMs in FM conditions; control CMs (FM vs. GM conditions). Significant genes (P<0.05, Wald test) are marked with “*”. B. Gene expression of acid ceramidase *ASAH1* from RNAseq analysis C. Heatmap showing log-fold changes in gene transcripts encoding lipid oxidation and lipid uptake components.

